# Structure and mechanism of TagA, a novel membrane-associated glycosyltransferase that produces wall teichoic acids in pathogenic bacteria

**DOI:** 10.1101/507483

**Authors:** Michele D. Kattke, Jason E. Gosschalk, Orlando E. Martinez, Garima Kumar, Robert T. Gale, Duilio Cascio, Michael R. Sawaya, Martin Philips, Eric D. Brown, Robert T. Clubb

## Abstract

*Staphylococcus aureus* and other bacterial pathogens affix wall teichoic acids (WTAs) to their surface. These highly abundant anionic glycopolymers have critical functions in bacterial physiology and their susceptibility to β-lactam antibiotics. The membrane-associated TagA glycosyltranserase (GT) catalyzes the first-committed step in WTA biosynthesis and is a founding member of the WecB/TagA/CpsF GT family, more than 6,000 enzymes that synthesize a range of extracellular polysaccharides through a poorly understood mechanism. Crystal structures of TagA from *T. italicus* in its apo- and UDP-bound states reveal a novel GT fold, and coupled with biochemical and cellular data define the mechanism of catalysis. We propose that enzyme activity is regulated by interactions with the bilayer, which trigger a structural change that facilitates proper active site formation and recognition of the enzyme’s lipid-linked substrate. These findings inform upon the molecular basis of WecB/TagA/CpsF activity and could guide the development of new anti-microbial drugs.

**AUTHOR SUMMARY:** Gram-positive bacteria cause thousands of deaths in the United States each year and are a growing health concern because many bacterial strains have become resistant to commonly used antibiotics. One of the most abundant polymers displayed on the surface of Gram-positive bacteria is wall teichoic acid (WTA), a negatively charged carbohydrate polymer that has critical functions in cell division, morphology, adhesion and pathogenesis. The WTA biosynthetic pathway has drawn significant interest as a drug target because clinically important methicillin-resistant *S. aureus* (MRSA) strains that lack WTA are defective in host colonization and re-sensitized to β-lactam antibiotics. To understand how bacteria produce WTA, we determined the structure and deduced the enzymatic mechanism of TagA, an important enzyme that is required for WTA synthesis. This research reveals a new method for enzyme regulation, whereby peripheral membrane association enables TagA to adopt its active form as a monomer. As TagA enzymes are highly conserved in bacteria, they can be expected to operate through a similar mechanism. The results of this work provide insight into WTA biosynthesis and could lead to innovative approaches to treat infections caused by pathogenic bacteria.

## INTRODUCTION

The thick peptidoglycan (PG) sacculus that surrounds Gram-positive bacteria maintains cellular integrity and is affixed with proteins and glycopolymers that have important roles in microbial physiology and host-pathogen interactions. In *Staphylococcus aureus* and other Gram-positive bacteria, wall teichoic acids (WTAs) are a major component of the cell wall, constituting up to 60% of its dry mass (1). WTAs have essential functions, including regulating PG biosynthesis, morphogenesis, autolysin activity, immune evasion, resistance to host cationic antimicrobial peptides, and pathogenesis (2–9). The WTA biosynthetic pathway has drawn considerable interest as an antibiotic target, as genetically eliminating WTA production in clinically important Methicillin-resistant *Staphylococcus aureus* (MRSA) re-sensitizes it to β-lactam antibiotics and attenuates its virulence (2, 3).

WTA polymers are constructed from polymerized alditol-phosphate subunits that are attached to the cell wall via a disaccharide-containing linkage unit. While the chemical structure of the main chain of the polymer can vary, the structure of the linkage unit is highly conserved across different species of Gram-positive bacteria and is composed of an *N-*acetylmannosamine (ManNAc) (β1→4) *N*-acetylglucosamine (GlcNAc) disaccharide appended to one to three glycerol-3-phosphate (GroP) groups (10) (**Fig. 1A**). The linkage unit performs a key function in WTA display, connecting the WTA polymer to the C6 hydroxyl of PG’s *N-*acetylmuramic acid (MurNAc) (11). WTA is synthesized on the cytoplasmic face of the cell membrane by modifying a membrane-embedded undecaprenyl-phosphate (C_55_-P) carrier. In *Bacillus subtilis* and *S. aureus,* the conserved GlcNAc-ManNAc-GroP linkage unit is first synthesized by the sequential action of the TagO, TagA, and TagB enzymes (originally designated TarOAB in *S. aureus*). TagO initiates WTA synthesis by transferring GlcNAc from the UDP-activated sugar to the C_55_-P carrier to produce lipid-α (12). The TagA glycosyltransferase (GT) then appends ManNAc from a UDP-ManNAc donor, producing a C_55_-PP-GlcNAc-ManNAc disaccharide-lipid product (lipid-β) (13, 14).

**Figure 1:**
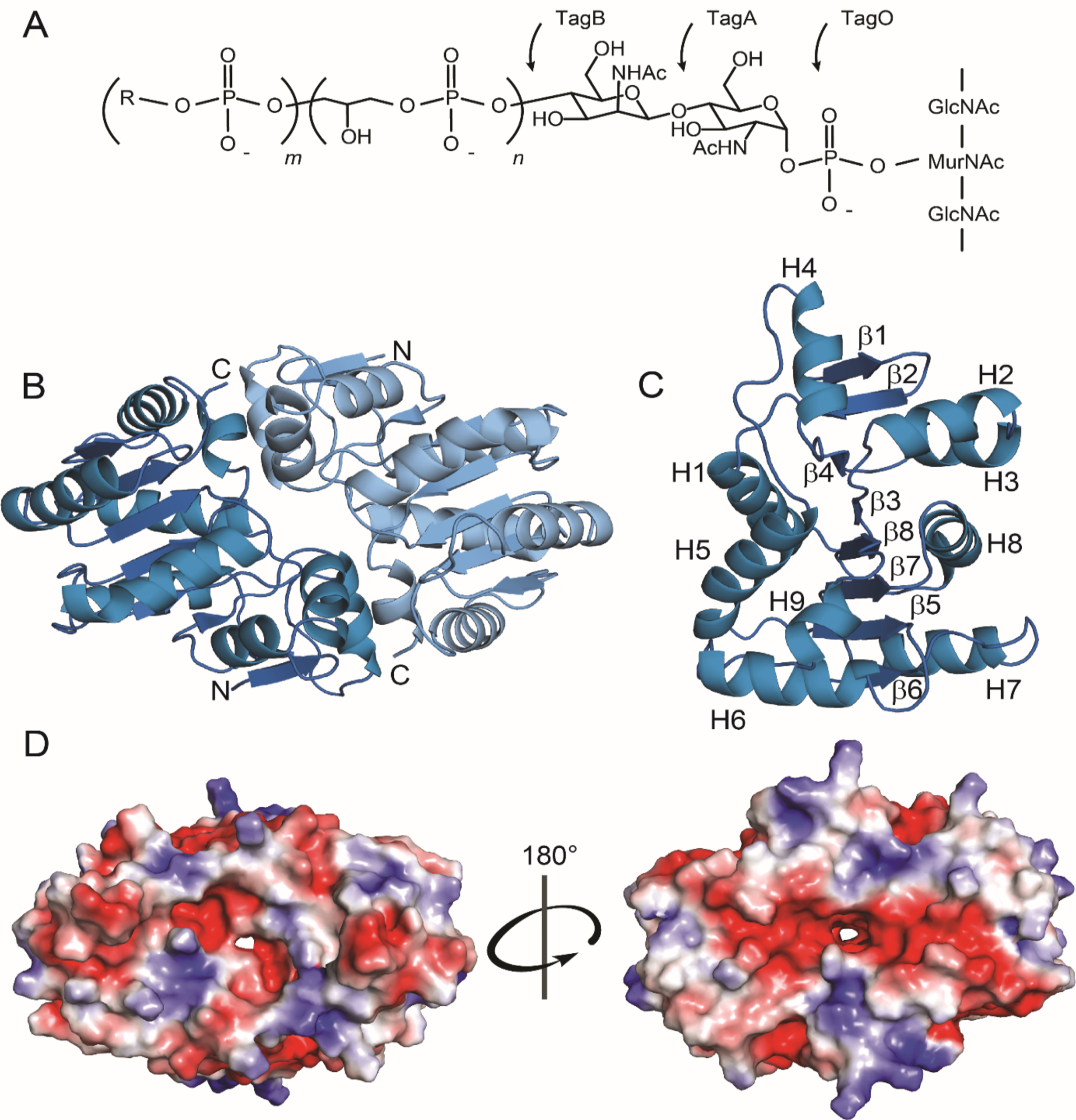
Kattke and Gosschalk *et al*. The WTA linkage unit and TagA structural characteristics. (A) WTA linkage unit and polymer. The linkage unit is attached to the peptidoglycan via the C6-hydroxyl of *N*-acetylmuramic acid and is composed of a GlcNAc (TagO-catalyzed), ManNAc (TagA-catalyzed), and *n =* 2-3 glycerol phosphates (TagB- and TagF-catalyzed). R = glycerol or ribitol, *m =* 40-60. (B) Cartoon ribbon representation of TagA^ΔC^ from *T. italicus.* Apo-TagA^ΔC^ crystallizes as a dimer. The dimer interface is over 1000 Å in surface area and is formed by buried hydrophobic residues. (C) TagA^ΔC^ protomer with secondary structural elements indicated. H = alpha helix, β = beta strand. (D) Electrostatic surface representation of the TagA^ΔC^ dimer. Negatively charged (*red*), neutral (*white*), and positively charged (*blue*) residues are indicated. Rotation of 180° about the dimer interface allows visualization of the pore.

Linkage unit synthesis is then completed by TagB, which appends a single GroP to lipid-β using a CDP-glycerol substrate that is produced by the TagD enzyme (13). In *S. aureus,* the linkage unit is then primed by TarF, which attaches at least one additional GroP. The TarIJL enzymes then construct the main chain of the polymer by adding 40-60 ribitol-5-phosphate (RboP) units (15–17). After being modified with GlcNAc by TarM and TarS, the TarGH ABC-like transporter exports the polymer to the cell surface, where it is further modified with D-alanine to tune its electrostatic properties (18). The assembled polymer is then covalently attached to the cell wall by an LCP ligase, which catalyzes a phosphotransfer reaction that joins WTA via its linkage unit to PG’s MurNAc (19–21). *B. subtilis* also uses functionally analogous enzymes to produce strain-specific GroP (spp. 168) or RboP (spp. W23) WTA polymers. Recent structural studies have begun to reveal the mechanism through which bacteria produce WTA, including how the polymerization is primed (TarF), and how WTA is modified with GlcNAc (TarM and TarS) and attached to the cell wall (LCP ligases) (22–24). However, it remains unknown exactly how bacteria produce the highly-conserved linkage unit that connects WTA to the cell wall.

The TagA N-acetylmannosamine transferase catalyzes the first committed step in WTA biosynthesis. It is an attractive target for new therapeutics aimed at treating MRSA infections, as *tagA-* strains are attenuated in virulence and re-sensitized to methicillin, imipenem, and ceftazidime. (25–28). TagA is also a founding member of the WecB/TagA/CpsF family of GTs (PFAM03808; CAZy GT26), which has over 6,000 members (29, 30). In addition to WTA, these enzymes synthesize a range of important surface-associated and secreted glycopolymers that function as virulence factors, including capsular polysaccharides of Group B Streptococcus (GBS) and the enterobacterial common antigen present in the outer-membrane of *Escherichia coli* and other Gram-negative bacteria (31–33). Industrially, this family is important as it includes the GumM GT, an essential enzyme in xanthan gum synthesis in *Xanthomonas campestris* (34). WecB/TagA/CpsF GTs are distinguished by their ability to elaborate membrane-embedded polyprenol substrates, but the molecular basis of their function remains unknown. Here, we report the crystal structure and biochemical studies of TagA from *Thermoanaerobacter italicus*, a close homolog of *S. aureus* TagA. Our results reveal that WecB/TagA/CpsF enzymes adopt a unique GT-E fold and shed considerable light onto their mechanism of catalysis. We propose that membrane association activates TagA by triggering a unique dimer to monomer quaternary structural change that facilitates lipid-α recognition and the formation of a catalytically competent active site. The results of these structural and mechanistic studies represent a major advancement in our understanding of WTA biosynthesis that could facilitate the discovery of new antibiotics that work by disrupting the synthesis of this important bacterial surface polymer.

## RESULTS AND DISCUSSION

### WecB/TagA/CpsF Enzymes are Structurally Novel Glycosyltransferases

To gain insight into the mechanism of catalysis, we determined the 2.0 Å crystal structure of the TagA enzyme from *Thermoanaerobacter italicus* (TagA^ΔC^, residues Met1-Gly195) (**Fig. 1b**). TagA^ΔC^ contains the highly conserved amino acid region that defines the WecB/TagA/CpsF family (**Fig. S1**), but lacks 49 C-terminal residues that target the protein to the membrane (*vide infra*). TagA^ΔC^ is much better suited for structural analyses, since unlike the full-length protein, it does not require high concentrations of salt and glycerol to be solubilized. Selenomethionine (SeMet)-labeled TagA^ΔC^ in its apo-form crystallized in the P2_1_ space group as a dimer, with eight molecules per asymmetric unit. The structure was determined using the multiple anomalous dispersion (MAD) method and is well-defined by continuous electron density. The subunits in the dimer are related by two-fold non-crystallographic symmetry and possess similar atomic structures; their heavy atom coordinates can be superimposed with an RMSD of 0.11 Å. Complete data collection and structural statistics are provided in **Table 1**.

**Table 1:**
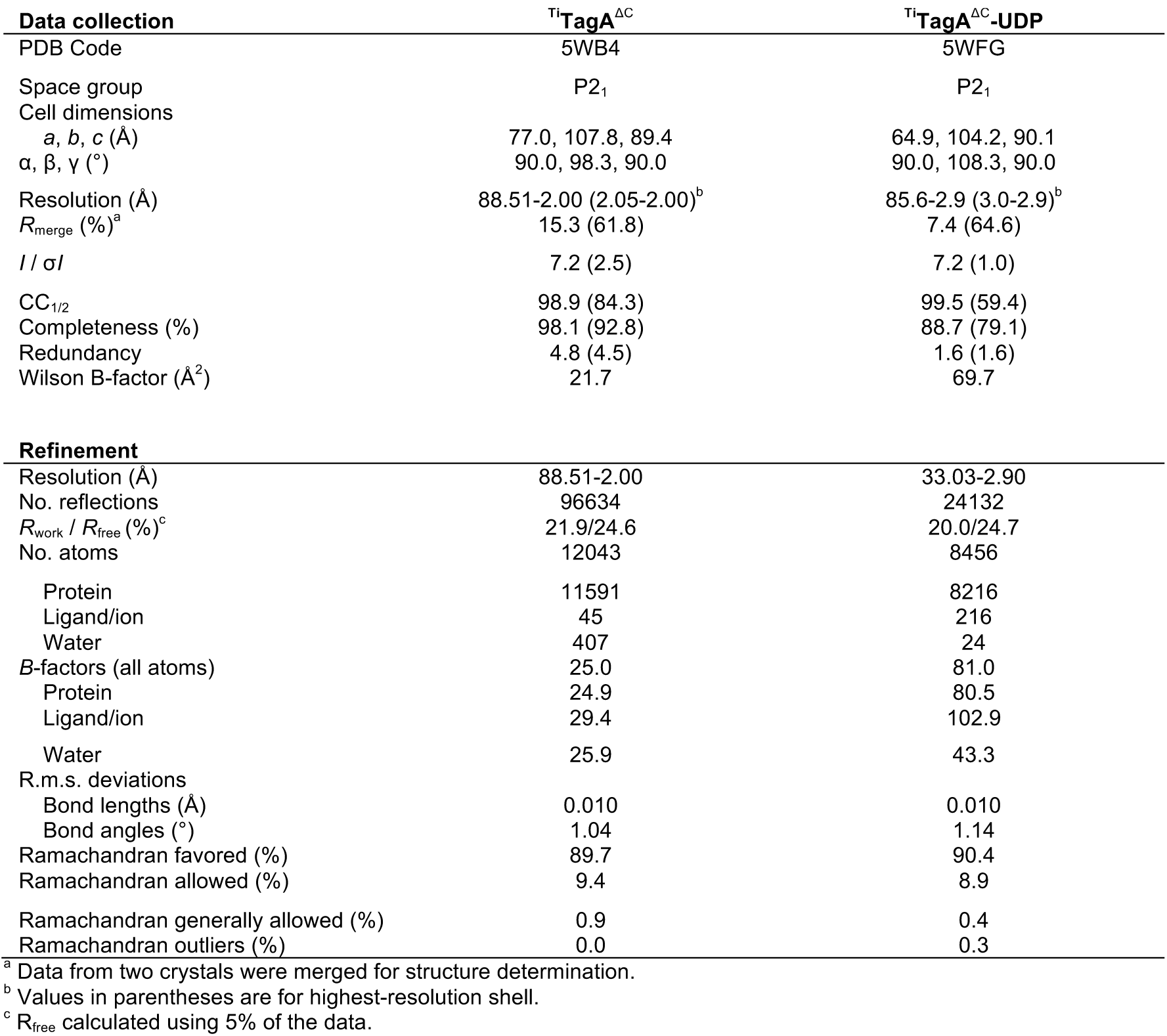
*Crystal data collection and structure refinement statistics.*

TagA adopts a unique α*/*β tertiary structure that differs markedly from previously described GTs (35, 36). Each protomer consists of eight β—strands and nine α-helices that form two distinct regions (**Fig. 1c**). The N-terminal region of TagA is formed by helices H2 to H4 that pack against a β-hairpin constructed from strands β1 and β2, while its larger C-terminal region is comprised of a six-stranded parallel β-sheet (β—strands β4, β3, β8, β7, β5, and β6) that is surrounded by seven helices (helices H1, H3, and H5 to H9). The arrangement of the six parallel strands forming the β—sheet resembles a Rossmann fold commonly found in nucleotide-binding proteins. The regions are interconnected, with the N-terminal β-hairpin forming a single backbone-backbone hydrogen bond to strand β4 within the C-terminal region (between the amide of Asp13 (β2) and the carbonyl of Asn63 (β3)). In the dimer, the C-terminal helix H8 in one subunit packs against helices H2 and H4 located in the N-terminal region of the adjacent protomer, burying 1,226 Å^2^ of solvent accessible surface area to produce a narrow pore (**Fig. 1d**). Analytical ultracentrifugation (AUC) experiments indicate that TagA^ΔC^ dimerizes in solution with relatively weak affinity, with a monomer-dimer dissociation constant (K_d_) of only 7.4 ± 0.7 μM (**Fig. S3A)**. As WecB/TagA/CpsF enzymes exhibit related primary sequences, it is expected that they will adopt tertiary structures that are similar to that observed for TagA.

Glycosylation reactions catalyzed by GT enzymes play a central role in biology, creating an enormous array of biologically important oligosaccharides and glycoconjugates. Interestingly, the array of enzymatic machinery used to perform glycosylation is surprisingly simple, and only four distinct GT protein folds have been identified that are capable of glycosyltransferase activity (termed GT-A, GT-B, GT-C and GT-D enzymes) (32, 36, 37). TagA^ΔC^ differs markedly from all of these enzymes based on its tertiary structure and how its secondary structural elements are arranged. Notably, TagA lacks the canonical Asp-X-Asp motif found in GT-A enzymes that participate in nucleotide binding and it differs substantially from GT-B and GT-C class enzymes that adopt multi-domain structures (32, 36). Interestingly, TagA does exhibit limited structural homology with DUF1792 (PDB ID: 4PFX), the founding member of the GT-D family that transfers glucose from UDP-glucose onto protein O-linked hexasaccharides (37). The backbone coordinates of a subset of residues within the TagA^ΔC^ and DUF1792 structures can be superimposed with an RMSD of 3.7 Å (**Fig. S2A**). However, consistent with these enzymes sharing only 15% sequence identity, the arrangement, number, and topology of their secondary structural elements are distinct (**Fig. S2B**). Furthermore, the enzymes have different catalytic mechanisms, as TagA exhibits ion-independent glycosyltransferase activity and it lacks the conserved Asp-X-Glu motif present in DUF1792 that coordinates an Mn^2+^ ion cofactor (14). Thus, TagA and related members of the WecB/TagA/CpsF family adopt a novel glycosyltransferase fold, which we term GT-E.

### Active site architecture

To define the enzyme active site, we determined the structure of the TagA^ΔC^:UDP complex at 3.1 Å resolution (**Fig. 2a**). The crystal structure visualizes the enzyme-product complex, as steady-state kinetics studies of *B. subtilis* TagA have shown that the enzyme operates via an ordered Bi-Bi mechanism in which UDP-ManNAc binds first and UDP is released last (14). The complex crystallizes as a dimer of trimers in the P2_1_ space group, with six molecules in each asymmetric unit (**Fig. S4**). The coordinates of the apo- and UDP-bound forms of TagA^ΔC^ are nearly identical (RMSD of 0.18 Å), suggesting that the enzyme binds UDP through a lock-and-key mechanism. In each protomer, UDP contacts the β7-H8 motif within the Rossmann-like fold, as well as the C-terminal edge of strand β5 (**Fig. 2a**). The uracil base is engaged in pi stacking with Tyr137 (β6-H7 loop), while Asp191 (H9) contacts the ribose sugar. The trimeric oligomer observed in the structure of the complex is presumably an artifact of crystallization, as AUC experiments performed in the presence of saturating amounts of UDP indicate that similar to apo-TagA^ΔC^, the UDP-bound protein fits best to a monomer-dimer equilibrium model rather than monomer-trimer or dimer-trimer equilibrium models (K_d_ = 3.5 ± 0.4 μM in the presence of 10:1 UDP:TagA) (**Fig. S3B**). In the complex, each UDP molecule is primarily contacted by a single subunit. However, the C2 hydroxyl group in each UDP molecule forms a hydrogen bond with the backbone carbonyl of Val192 (H9) located in the adjacent subunit, which may explain why the complex crystallized as a trimer.

**Figure 2:**
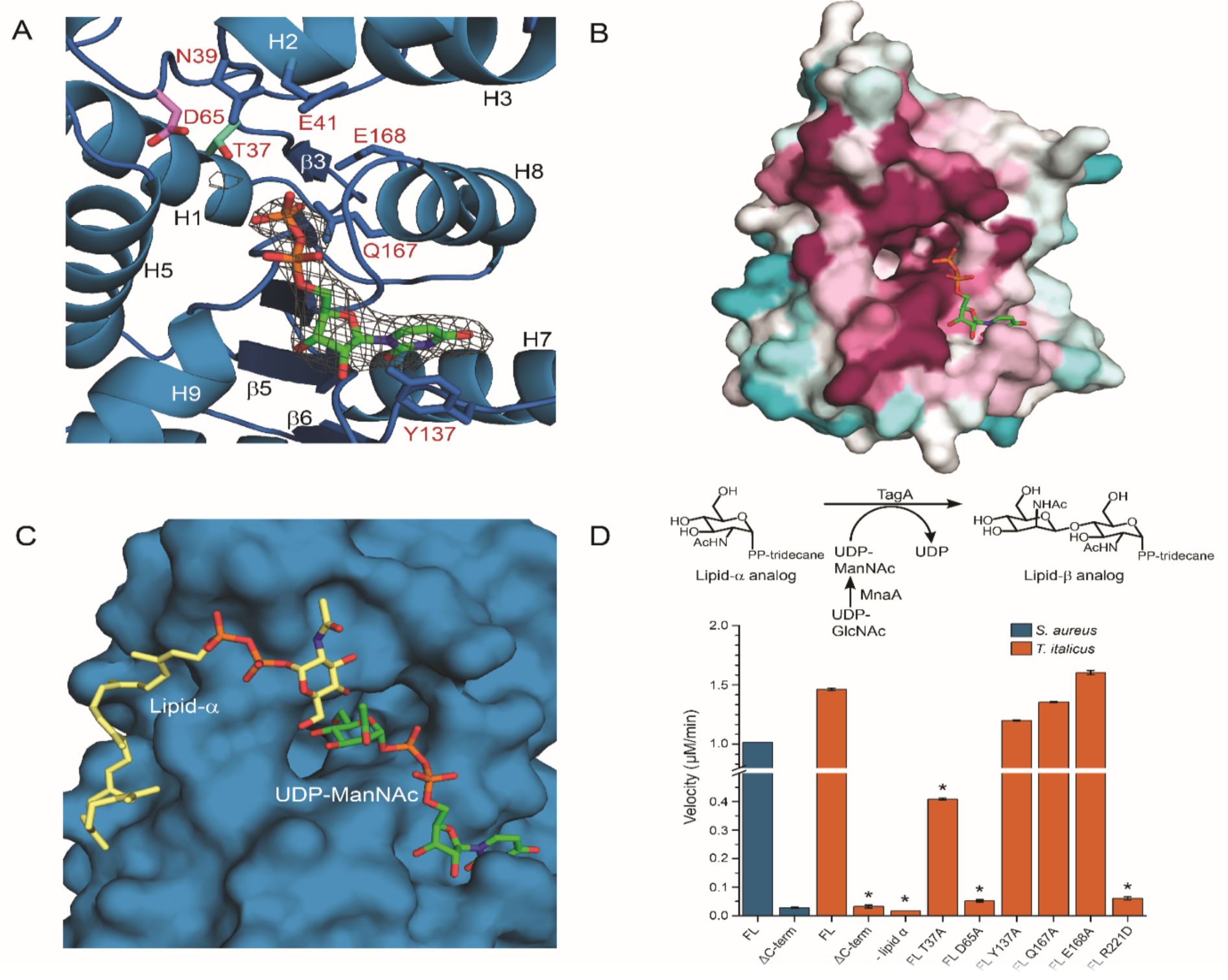
Kattke and Gosschalk *et al*. Substrate binding and mechanistic studies of TagA. (A) The electron density of UDP in the UDP:TagA^ΔC^ crystal structure places the β-phosphate of UDP (*orange*) adjacent to the proposed catalytic base, Asp65 (*pink*), and putative ManNAc stabilizing residue, Thr37 (*cyan*). The uracil nucleoside is pi-stacked over Tyr137. (B) Consurf analysis reveals that UDP projects its phosphates (*orange*) into the pocket and is coordinated by a pocket of highly conserved residues. Highly conserved (*magenta*), moderately conserved (*white*) and weakly conserved (*teal*) residues are indicated. (C) Proposed enzyme-substrate complex of the TagA protomer generated by Autodock vina with a lipid-α analog (*yellow*) and UDP-ManNAc (*green*). The coordinates of UDP from the UDP:TagA^ΔC^ trimer crystal structure were used to restrain UDP-ManNAc, and the docking results position the non-reducing end of GlcNAc toward the C4 in ManNAc when bound to the TagA^ΔC^ dimer. (D) The upper panel indicates the reaction scheme used for the *in vitro* TagA activity assay. 200 nM TagA enzyme is incubated at 30°C with 100 µM lipid-α substrate analog and UDP-ManNAc produced *in situ* from UDP-GlcNAc by the epimerase MnaA, followed by quenching with 4M urea. Conversion of UDP-ManNAc to UDP is monitored at 271 nm using a DNAPak PA200 anion exchange column. The lower panel indicates activity measurements of *T. italicus* and *S. aureus* TagA enzymes from the *in vitro* TagA activity assay, as described above. Reactions with error bars were performed in triplicate, and asterisks indicate p<0.005 by Student’s T-test.

Intriguingly, positioned immediately adjacent to the UDP binding site on each protomer is a large pocket that harbors several phylogenetically conserved amino acids that are important for catalysis (**Fig. 2b**). The pocket resides near the C-terminal end of the parallel β3 and β8 strands and has walls that are formed by residues located in helices H5 and H8, as well as residues within the polypeptide segments that connect strand β7 to helix H8, and strand β3 to helix H2. Several highly-conserved residues are located within the pocket and its periphery, including Thr37, Asn39, Asp65, Arg83, Gln167, and Glu168 (**Figs. 2a** and **S1**). Interestingly, in the crystal structure of the TagA^ΔC^:UDP complex, the β-phosphate group of UDP extends inward toward the pocket. As UDP is a competitive inhibitor of the UDP-ManNAc substrate, it is likely that these ligands bind to the same site on the enzyme, such that the ManNAc moiety within UDP-ManNAc is projected into the pocket where it can interact with the conserved side chains of residues Thr37, Gln167 or Glu168 (14). Modeling studies of substrate binding to the TagA^ΔC^ protomer using Autodock vina (see *Materials and Methods*) suggest that the UDP-ManNAc and lipid-α substrates bind to opposite sides of the conserved pocket (38). In models of the enzyme-substrate ternary complex, the GlcNAc and diphosphate portion of lipid-α are positioned near residues that connect strands β4 to helix H4, while the undecaprenyl chain of lipid-α exits near the C-terminus of the TagA^ΔC^ protomer (**Fig. 2c**). In this binding mode, the sugar acceptor’s C-4 hydroxyl group is positioned near the side chain of Asp65, while the highly-conserved side chains of Arg83 and Asn39 are adjacent to lipid-α’s diphosphate and GlcNAc, respectively. To investigate the importance of these conserved pocket residues, we reconstituted its GT activity *in vitro* using UDP-ManNAc and a lipid-α analog that replaces its undecaprenyl chain with tridecane, as previously reported (14, 39). The full-length TagA enzymes from *T. italicus* and *S. aureus* exhibit similar transferase activities *in vitro*, producing the UDP product at a rate of 1.5 *µ*M min^-1^ and 1.0 *µ*M min^-1^ at 200 nM enzyme concentration, respectively (**Fig. 2d**). This is expected, as all TagA homologs presumably catalyze the synthesis of the ManNAc(β1→4)GlcNac glycosidic bond within the conserved linkage unit. Thr37Ala and Asp65Ala mutations in TagA cause the largest decreases in activity (0.41 *µ*M min^-1^ and 0.052 *µ*M min^-1^, respectively), compatible with these residues residing within the enzyme’s active site. As discussed later, the carboxyl side chain of Asp65 is poised to function as general base that deprotonates the nucleophilic C4 hydroxyl group in GlcNAc, while the side chain of Thr37 may stabilize the orientation of ManNAc by interacting with its C5 hydroxyl group.

### A conserved C-terminal appendage is required for catalysis

Surprisingly, the mutational analysis reveals that only the full length TagA protein is enzymatically active *in vitro*, while the truncated TagA^ΔC^ protein used for crystallography and modeling is catalytically inactive (**Fig. 2d**). This is compatible with the high level of primary sequence conservation of C-terminal residues within WecB/TagA/CpsF enzymes and suggests that the deleted appendage may be necessary to construct a catalytically competent active site (**Fig. S1**). To gain insight into the function of the appendage, we utilized the structure of the TagA protein modeled using Generative Regularized Models of Proteins (GREMLIN), a recently developed protein modeling server that predicts tertiary structure by exploiting sequence conservation and amino acid co-evolutionary patterns (40, 41). Only GREMLIN-predicted structures in which TagA was assumed to be monomeric yielded favorable results, and is substantiated by important correlations between residues within the body of TagA^ΔC^ and the appendage (in *T. italicus:* Arg83 and Arg205, Ala72 and Val235, Gln47 and Lys234, Val197 and Glu218/Lys217, and Lys198 and Lys217 couplings). In general, the tertiary structures of the GREMLIN-predicted model of monomeric TagA (TagA^GM^) and the experimentally determined crystal structures of TagA^ΔC^ are similar. However, only in TagA^GM^ is the C-terminal appendage present, which forms three α—helices (H10 to H12) that pack against the active site harboring the catalytically important Asp65 and Thr37 residues. The C-terminal appendage is presumably unstructured in dimeric forms of the enzyme, as, importantly, the contact surface used to engage the appendage is occluded by inter-subunit interactions in the crystal structures of TagA^ΔC^.

The TagA^GM^ monomer provides insight into why the C-terminal appendage is critical for catalysis, as it contains several highly-conserved arginine residues that project side chains into the enzyme’s active site (Arg214, Arg221 and Arg224) (**Fig. 3a**). Of particular interest is the side chain of Arg221 within helix H11 of the appendage, as modeling of TagA^GM^ bound to its substrates suggests that the Arg221 guanidino group may stabilize the β-phosphate of the UDP leaving group (**Fig. 4a**). Indeed, Arg221Glu mutation in TagA significantly abates TagA GT activity, strongly suggesting that only the monomeric form of TagA is enzymatically active (**Fig. 2d**). While not tested in this study, the side chains of Arg214 and Arg224 may also be important for catalysis, as some cation-independent GTs use more than one positively charged residue to stabilize phosphotransfer reaction intermediates (42). Chemical crosslinking studies of cells expressing full length *T. italicus* TagA indicate that the protein forms a mixture of monomers and dimers **(Fig. S3C)**, consistent with the relatively weak dimerization affinity of TagA^ΔC^ measured by AUC (**Fig S3A** and **S3B**). Thus, in the cell, TagA presumably toggles between distinct oligomeric states: 1) enzymatically active monomers, in which the C-terminal appendage contributes key residues to the active site, and 2) inactive dimers, in which inter-subunit interactions mask an incompletely formed active site.

**Figure 3:**
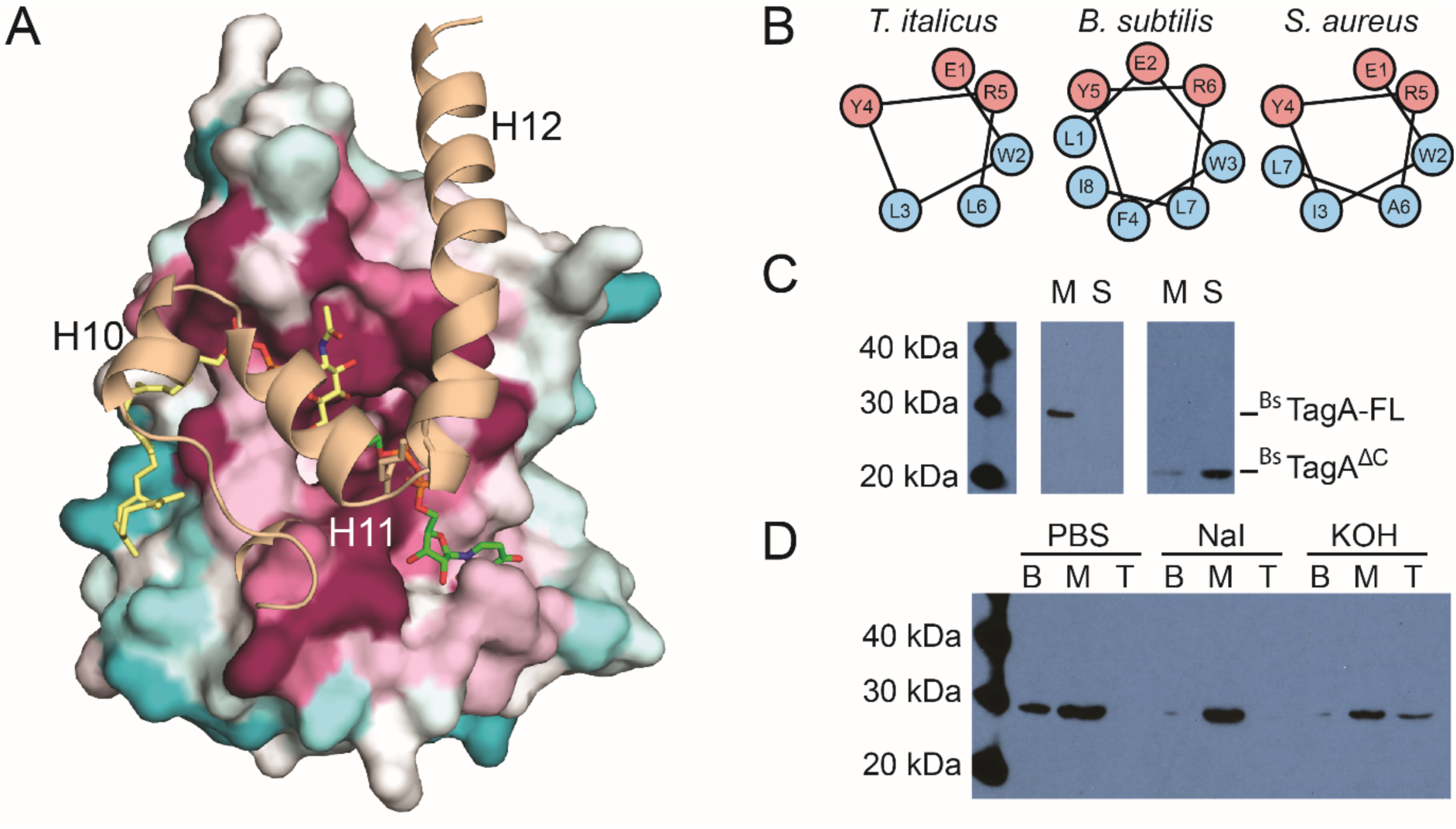
Kattke and Gosschalk *et al*. Computational and biochemical studies of the TagA enzyme inform cellular localization. (A) A model of full-length TagA was constructed with experimentally determined TagA^ΔC^ (*surface representation*) and the C-terminal domain, which was modeled by GREMLIN structural prediction (*cartoon representation*). Three C-terminal helices appear to complete the active site and obstruct the dimeric interface of TagA^ΔC^. (B) Helical wheel projections of helix H11 in TagA homologs predict a putative amphipathic helix. (C) TagA associates with the bacterial cell membrane. Immunoblots of cellular fractionation indicate that *B. subtilis* TagA is exclusively localized to the membrane (*M*), while TagA^ΔC^ is primarily localized in the supernatant (*S*). Samples were fractionated by ultracentrifugation identically and the ^Bs^TagA-FL blot was exposed for 10 minutes, while the ^Bs^TagA-V196 blot was exposed for 1 minute. (D) TagA is a peripheral membrane protein. Chaotropic and alkaline treatments of *B. subtilis* TagA reveal that the enzyme is peripherally associated with the membrane and is more effectively displaced by alkaline treatment. Treated membrane fractions were loaded onto a sucrose cushion, centrifuged, and carefully separated into bottom (*B; pellet*), middle (*M; sucrose cushion volume*), and top (*T; sample volume*).

**Figure 4:**
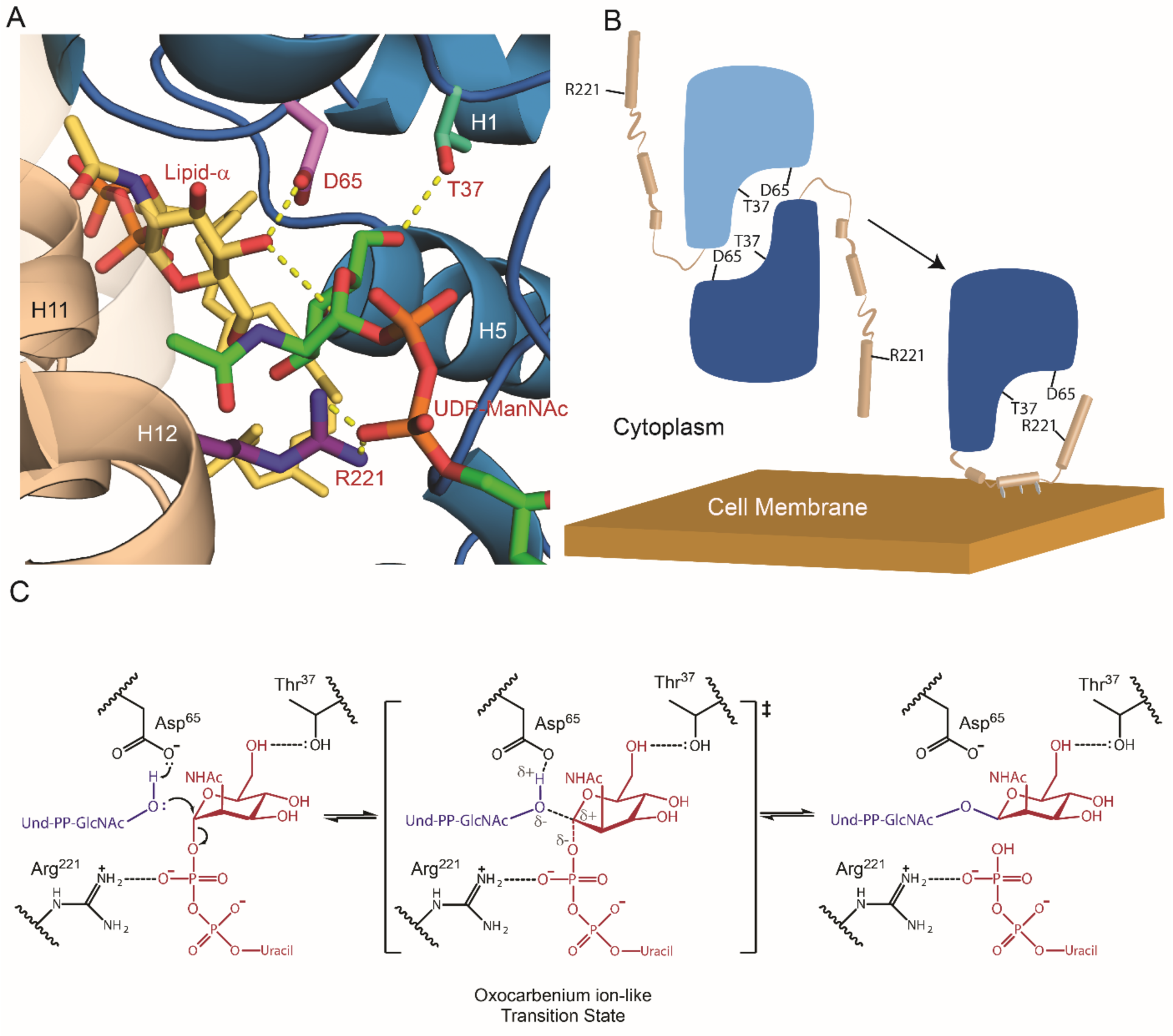
Kattke and Gosschalk *et al*. Model of the TagA enzyme-substrate complex and mechanism of catalysis. (A) The proposed active site of TagA co-localizes residues D65, T37, and R221. Lipid-α (*yellow*) is activated by the catalytic base D65 (*pink*), while ManNAc (*green*) is positioned by contacts between its C6 hydroxyl and T37 (*cyan*). The C-terminal helices (*tan*) are modeled according to GREMLIN structural predictions and place R221 (*purple*) adjacent to the phosphates of UDP-ManNAc (*orange*) to putatively stabilize the leaving group. (B) The TagA molecular mechanism is proposed to utilize a dimer to monomer transition to regulate glycosyltransferase activity. TagA is stabilized as a soluble dimer. Upon interaction with the cell membrane, the C-terminus adopts an ordered state and disrupts the dimer interface, which produces a competent active site by co-localizing D65, T37, and R221 to coordinate the soluble UDP-ManNAc and membrane-bound lipid-α substrates. (C) TagA reveals the catalytic mechanism of the GT26 family. Asp65 activates lipid-α, which proceeds to attack UDP-ManNAc in an S_N_2-like mechanism. Coordination between Arg221 and the phosphates of UDP stabilize the leaving group, permitting the oxocarbenium ion-like transition state. The mechanism is completed by glycosidic bond-formation between GlcNAc and ManNAc to form lipid-β.

### Membrane-induced structural changes likely facilitate the recognition of bilayer-embedded polyprenol substrates

To build the linkage unit, TagA should associate with the cytoplasmic membrane where it attaches ManNAc to its lipid-α substrate. Intriguingly, only the monomeric form of TagA contains a non-polar surface patch that is suitable for interacting with the lipid bilayer. In monomeric TagA^GM^, helices H11 and H12 within the C-terminal appendage project several non-polar side chains into the solvent for potential membrane interactions (e.g. Leu212, Ile216, Ile233, in *T. italicus* TagA) (**Fig. 3b**) (43–47). To investigate the role of the C-terminal appendage in membrane binding, we determined how this structural element affected TagA localization in *B. subtilis*, since unlike *T. italicus,* robust tools are available to genetically manipulate this model Gram-positive bacterium. Importantly, the TagA homologs in these organisms are related (34% sequence identity) and both contain the conserved C-terminal appendage (**Fig. S1**). *B. subtilis* cells expressing hexahistidine-tagged TagA (^Bs^TagA) were fractionated and analyzed by Western blotting. ^Bs^TagA peripherally associates with the membrane, since it is present in the membrane fraction, but released into the soluble fraction after adding either potassium hydroxide or the chaotropic salt sodium iodide (**Fig. 3c** and **3d**, respectively). Interestingly, TagA lacking the C-terminal appendage (^Bs^TagA^ΔC^, residues Met1-Val196) primarily partitions into the soluble fraction, indicating that this structural element tethers the protein to the membrane (**Fig. 3c**). Other bacterial enzymes are targeted to the membrane via terminal helices (e.g. TagB, FtsA, MinD, PBP enzymes), but TagA is novel because its helices form an integral part of the active site (43–47).

Collectively, our data suggest that TagA is activated through a unique membrane-association mechanism that is mediated by residues at its C-terminus. TagA is in equilibrium between monomeric and dimeric states. Removed from the membrane, it is primarily dimeric with interfacial interactions obstructing the binding site for its C-terminal appendage and holding the enzyme in an inactive, dimeric state (**Fig. 4B**). However, upon encountering the membrane, we posit that non-polar side chains within the appendage are inserted into the bilayer, thereby nucleating its folding and the formation of a monomeric, catalytically functional enzyme in which the appendage contributes key active site residues. In the TagA^GM^ monomer, a gap is located between helices H11 and H12 in the C-terminal appendage, forming a short pore that connects the active site to the protein’s membrane binding surface. Thus, membrane induced folding of the C-terminal appendage may also facilitate recognition of the lipid-α by providing an additional binding site for several of its non-polar prenyl groups. While the catalytic activities of other membrane-associated enzymes are regulated by lipid bilayer-induced quaternary structural changes or allosteric mechanisms, to the best of our knowledge, the TagA activation mechanism outlined here is unique (48).

In conclusion, our results provide direct insight into how TagA enzymes synthesize the conserved linkage unit used to attach WTA to the cell wall and, and more generally, how members of the large WecB/TagA/CpsF GT family produce a range of important surface-associated and secreted bacterial glycopolymers. TagA is a structurally unique GT that we propose defines a new GT-E fold. Removed from the membrane, it forms catalytically inactive dimers that are presumably incapable of mediating spurious GT reactions or hydrolyzing UDP-ManNAc. However, upon encountering the membrane containing its lipid-α substrate, conserved C-terminal residues in TagA fold into an essential active site appendage, stabilizing the monomeric and catalytically-active form of the enzyme. From our structural and biochemical data, a working model of the catalytic mechanism can be proposed (**Fig. 4C**). Previous studies have shown that catalysis likely occurs via an S_N_2-like mechanism that inverts the anomeric stereochemistry of ManNAc (14). It seems likely that Asp65 functions as a base, deprotonating the GlcNAc C4 hydroxyl in lipid-α. This would facilitate its nucleophilic attack at the anomeric carbon of ManNAc, resulting in an oxocarbenium ion-like transition state that is stabilized by electrostatic interactions between Arg221, donated by the C-terminal appendage, and the diphosphate moiety of UDP. Thr37 within the conserved pocket may play an important role in orienting the ManNAc moiety of the sugar donor, poising its electrophilic center for glycosidic bond formation. A complete understanding of the catalytic mechanism will require the structure determination of additional key reaction intermediates. As WTA and other bacterial glycopolymers are critical components of the cell wall, the results of these studies are of fundamental importance and could facilitate the discovery of GT-E (WecB/TagA/CpsF) specific enzyme inhibitors that could be useful antibiotics.

## MATERIALS AND METHODS

### Cloning, expression, protein purification, and crystallization

Bacterial strains and plasmids used in this study are listed in SI Materials and Methods, Table I. Protocols for protein purification, crystallization, and structure determination are detailed in SI Materials and Methods.

### Mutagenesis and activity assays

Conservation analysis was conducted using the Consurf Server (49–51). The lipid-α or UDP-ManNAc substrates were generated *in silico* using the Phenix electronic Ligand Builder and Optimization Workbench (Phenix.eLBOW) (52). Cis-trans configuration and stereochemistry were confirmed or corrected using the Phenix Restraints Editor Especially Ligands (Phenix.REEL). The substrates were docked to the 2.0 Å resolution protomer structure using Autodock vina with a 25 × 25 × 25 Å search space and an exhaustiveness of 18 (38). TagA *in vitro* enzyme activity was determined using an anion-exchange HPLC system and the following assay conditions: 0.2 µM TagA; 100 µM lipid-α; 100 µM UDP-GlcNAc; 3 µM MnaA; 50 mM Tris-HCl, pH 7.5; and 250 mM NaCl (39). Reactions were incubated for 40 minutes before quenching with 3 M urea. The reactions were separated with a DNAPak PA200 anion-exchange column using a buffer gradient of 100% Buffer A (20 mM NH4HCOO3, and 10% MeCN, pH 8.0) to 90% Buffer A plus 10% Buffer B (20 mM NH4HCOO3, 10% MeCN, and 1 M NaCl, pH 8.0) over 10 minutes. UDP-ManNAc or UDP elution peaks were monitored at 271 nm and integrated to determine turnover rate.

### Cell fractionation

Overnight *B. subtilis* cultures containing selective antibiotics were diluted 1:100 into 1 L of fresh LB broth containing 1 mM IPTG. Cultures were incubated at 37°C and 250 rpm until an OD_600_ of 1.0 was reached. Cells were pelleted at room temperature, washed once with PBS, and then frozen at −80°C. Cells were fractioned as previously reported with several exceptions (47). Cells were re-suspended in 10 mL of Lysis Buffer (PBS, pH 7.3; 1 mM EDTA; 1 mM DTT; 10 µg/mL RNase; 10 µg/mL DNase; 2 mM PMSF, and 100 µL Protease inhibitor cocktail) and sonicated on ice for eight minutes, with one minute “on” (one second pulses for 60 seconds) and one minute “off” (no pulses for 60 seconds). Ten milliliters of lysate were divided into two 5 mL volumes and centrifuged in a Beckman type 50 TI rotor in a Beckman XPN-100 preparative ultracentrifuge. The lysate was centrifuged at 9,600 rpm for 10 minutes to remove cellular debris. The supernatant was collected and centrifuged at 21,000 rpm. The supernatant was removed and spun for a third time at 40,000 rpm for one hour. This final supernatant fraction was considered to be the soluble fraction, and the remaining pellet, which contained the membrane fraction was re-suspended in 1 mL of ice-cold Lysis Buffer. The membrane fraction was diluted ten-fold for comparison with the supernatant and samples were run on an SDS-PAGE gel for 50 minutes at 170 V and transferred to a PVDF membrane using an iBlot transfer device (ThermoFisher Scientific). The membrane was fixed in methanol for five minutes, briefly washed in water, and then blocked over-day in TBST Blocking Buffer (20 mM Tris-HCl, pH 7.5; 500 mM NaCl; 0.05% Tween; and 5% w/v nonfat milk). The membrane was washed, incubated with primary antibody (Invitrogen #MA-21215, mouse anti-6_X_His, 1:1000 dilution in TBST Blocking Buffer) overnight at 4°C, washed again, and incubated with secondary antibody (Sigma-Aldrich #A9044, anti-mouse horseradish peroxidase). The membrane was incubated with SuperSignal West Pico Chemiluminescent Substrate (Thermo Scientific #34080) for five minutes and exposed to radiography film for one to ten minutes.

### Chaotropic agent analysis

Membrane fractions were prepared as described above and re-suspended to 600 µL. Fractionation was performed as previously reported, with some minor changes (53). Two-hundred microliters of membrane resuspension was diluted 1:3 in PBS containing 1.5 M sodium iodide, 0.1 N potassium hydroxide, or PBS alone. Samples were loaded onto a 2.4 mL sucrose cushion (PBS and chaotrope, 0.5 M sucrose) and centrifuged for 30 minutes at 40,000 rpm. The top (800 µL) and middle (2.4 mL) fractions were removed, and the pellet was re-suspended in 200 µL of PBS. The top and middle fractions were precipitated with 10% trichloroacetic acid on ice for 30 minutes, centrifuged at 20,000 *g* for 10 minutes, and then re-suspended in 200 uL of 8 M urea. Samples were mixed 1:1 with 2X SDS loading dye (100 mM Tris base, 200 mM DTT, 4% SDS, 0.2% bromophenol blue, 20% glycerol). Immunoblotting was performed as described above.

## ACKNOWLEDGMENTS

We thank members of the Brown and Clubb laboratories for critical review and discussion of the manuscript. We also thank Sergey Ovchinnikov from the Baker laboratory for discussion of the TagA model generated by GREMLIN structural prediction. We thank Dr. Beth Lazazzera for plasmids and her valuable insight. This research was supported by funding from the National Institutes of Health (AI52217 to R.T.C.). O.M. was supported by the UCLA Chemistry Biology Interface Training program (NIH T32GM008496). M.D.K. was supported by the UCLA-Molecular Biology Institute Whitcome Pre-Doctoral Training Grant. E.D.B. acknowledges support from a Foundation grant from the Canadian Institutes of Health Research (FDN-143215) and an award from the Canada Research Chairs program. R.T.G. was the recipient of a Canadian Institutes of Health Research graduate scholarship.

## Supplementary Information Text

### MATERIALS AND METHODS

#### Cloning, expression, and protein purification

The N-terminal domain of TagA from *T. italicus* (^Ti^TagA^ΔC^, residues Met1-Gly195) or *S. aureus* (^Sa^TagA^ΔC^, residues Ala10-Ala204) was expressed from a pMAPLe4 plasmid in *Escherichia coli* BL21(DE3) cells (Table S1). Standard methods were employed, with cultures grown in the presence of 50 µM kanamycin at 37°C until an OD_600_ of 0.6-0.8 was reached. Protein expression was initiated by adding isopropyl-β-D-1-thiogalactopyranoside (IPTG) to 1 mM, followed by overnight protein expression at 18°C. A four-liter cell culture was harvested by centrifugation at 7000 rpm in a Beckman JA-10 rotor, and the pellet was re-suspended in 40 mL of Buffer A (50 mM Tris, pH 7.5; 500 mM NaCl; 40 mM CHAPS) with 400 µL of protease inhibitor cocktail (Sigma) and 2 mM phenylmethanesulfonylfluoride (PMSF). The cells were then lysed using an EmulsiFlex high pressure homogenizer (Avestin). Cell lysates were fractionated by centrifugation at 15,000 rpm in a Beckman JA-20 rotor, and the soluble portion was applied to a gravity column containing 10 mL of suspended His-Pure Co^2+^ resin (Life Technologies) that was pre-equilibrated with Buffer A. The resin was washed with 20 mL aliquots of Buffer B (50 mM Tris, pH 7.5; 500 mM NaCl; 0.5% CHAPS) that contained 0, 25, or 50 mM imidazole. His-tagged TagA^ΔC^ was eluted using Buffer B with 500 mM imidazole, and the fractions were pooled and concentrated using a 10 kDa MWCO Amicon Ultra-15 centrifugal filter (Millipore). To remove His_6_-tag from the protein, TEV protease was added to TagA^ΔC^, and the solution was dialyzed in a 3.5 kDa MWCO Slide-A-Lyzer dialysis cassette (ThermoFisher Scientific) against Buffer C (50 mM Tris, pH 7.5; 200 mM NaCl) at 4 °C overnight. TEV protease was then separated from TagA^ΔC^ by binding 10 mL of suspended His-Pure Co^2+^ resin (Life Technologies) that was pre-equilibrated with Buffer C; cleaved TagA^ΔC^, which lacked the His_6_-tag, was eluted from the resin using Buffer C. Cleaved TagA^ΔC^ was further purified by gel filtration chromatography using a Sephacryl size-exclusion column (GE Healthcare Life Sciences) that was equilibrated with Buffer C. Purified TagA^ΔC^ was then pooled, concentrated to 55 mg/mL, and stored at 4°C.

Selenomethionine (SeMet) labeled protein was prepared with cultures grown in M9 minimal media in the presence of kanamycin at 37°C until an OD_600_ of 0.6-0.8 was reached. Protein expression was then initiated by adding IPTG to 1 mM followed by overnight protein expression at 18°C. Protein purification was performed as described for native protein.

#### Structure determination

Recombinant TagA^ΔC^ at a concentration of 50 mg/mL in Buffer C was used for crystal screening. Screening was performed with the JCSG+ broad matrix suite (Molecular Dimensions) at room temperature in a sitting-drop vapor diffusion format (200 nL drop size). SeMet-labeled protein crystals grew over the course of three days in the presence of 200 mM lithium sulfate; 100 mM phosphate citrate, pH 4.2; and 20% PEG 1000. For X-ray data collection, TagA^ΔC^ crystals were cryoprotected using reservoir solution containing 35% glycerol. Diffraction datasets were collected at the Advanced Photon Source (APS) beamline 24-1D-C equipped with a Pilatus-6M detector. All data were collected at 100 K. Data were collected at the detector distance of 300 mm, with 0.25° oscillations, and at a 0.9791 Å wavelength. Multi-wavelength anomalous dispersion (MAD) experiment was collected at peak (12663.0 eV), inflection (12660.3 eV), and high remote (12763.0 eV) energy wavelengths.

The TagA^ΔC^ crystals diffracted X-rays to 2.0 Å resolution. The XDS/XSCALE package was used to index, integrate and scale data in P2_1_ space group (1). The asymmetric unit of the crystal contained eight protein molecules, yielding a Matthews coefficient of 2.11 Å/Da and a 41.79% solvent content in the unit cell. The SHELX suite was used to locate the heavy atom substructure, which identified a total of 56 selenium atom sites (2). The quality of the phases calculated with the peak, inflection, and high remote energy diffraction datasets were improved using SHARP and the wARP suite (Global Phasing Limited). The heavy atom parameters were refined with MLPhare using the CCP4i suite (3). Density modification and non-crystallographic symmetry averaging was performed with the CCP4i suite to improve the quality of the electron density map. Automated model building was performed with BUCCANEER, followed by refinement with BUSTER (4). Modeling of the additional electron density was confirmed using 2F_o_-F_c_ omit maps generated using BUSTER. Complete refinement and structure statistics are reported in Table I (5WB4).

A second crystal form was produced with recombinant TagA^ΔC^ in the presence of UDP and ManNAc ligands. TagA^ΔC^ at a concentration of 45 mg/mL in Buffer C with 10 mM UDP and 10 mM ManNAc was used for crystal screening with the JCSG+ broad matrix suite (Molecular Dimensions) as described above. TagA^ΔC^-ligand co-crystals grew over the course of two days in the presence of 200 mM calcium acetate; 100 mM sodium cacodylate, pH 6.5; and 40% PEG 300. A single wavelength diffraction dataset for a non-cryoprotected crystal was collected at the APS beamline 24-1D-C equipped with a Pilatus-6M detector as described above. The crystals diffracted X-rays to 2.9 Å resolution. The XDS/XSCALE package was used to index, integrate and scale data in P2_1_ space group (1). The asymmetric unit of the crystal contained six protein molecules, yielding a Matthews coefficient of 2.22 Å/Da and a 44.66% solvent content in the crystal unit cell. The PHASER program in the CCP4i suite was used for molecular replacement, employing the coordinates of the apo-TagA^ΔC^ (5WB4) (5). Molecular replacement yielded a single solution, which was refined in iterative runs using Buster software. Additional electron density resembling the UDP ligand was observed using 2F_o_-F_c_ omit maps generated by BUSTER (4). Complete refinement and structure statistics are reported in Table 1 (5WFG).

TagA carboxyl terminal structure was modeled using Generative Regularized Models of Proteins (GREMLIN), accessed online at http://gremlin.bakerlab.org. TagA structure could not be modeled as a dimer in its full length (personal communications). Chain A of TagA^ΔC^ and TagA^GM^ were aligned with an RMSD of 2.2 Å.

#### Oligomeric Analysis

The dissociation constant and oligomeric determination of apo- and UDP-bound TagA^ΔC^ were determined by equilibrium sedimentation analytical ultracentrifugation (AUC) on an Optima XL-A analytical ultracentrifuge (Beckman Coulter, Brea, CA). Data regression analysis was performed using the Beckman-Coulter Optima Analytical Ultracentrifuge Origin Data Analysis Package. The data were fit to multi-exponential models and represented best by a monomer-dimer equilibrium that was calculated using the predicted monomeric molecular weight of 21633 Da by the ExPASy ProtParam tool. The dissociation constant (*K*_d_) was determined to be the inverse of *K*_a(conc)_ using equation 1 (6):

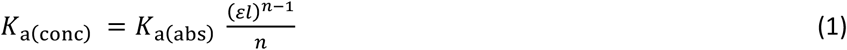

The molar extinction coefficient (*ε*) was determined to be 12950 cm^-1^M^-1^ using the ExPASy ProtParam tool, where *l* is the path length of 1.2 cm, *n* is the order of oligomerization, and *K*_a(abs)_ is the absorbance association constant, as determined by the nonlinear regression of the monomer-dimer multiexponential model using the analytical software program mentioned above. Chemical crosslinking experiments with disuccinyl suberate (DSS) (ThermoFisher Scientific) was performed according to manufacturer guidelines, with the following variations. RC21tagAFL and RC21tagAG195 overnight cultures were diluted 1:100 into LB, grown at 37°C and induced with 1mM IPTG at OD_600_ = 0.4. After two hours, cells were collected, washed three times with buffer (PBS, pH 8.0) and finally re-suspended at 3X concentration. Ten microliters of sample were loaded onto an SDS-PAGE gel, and western blotting was performed as described above.

#### *Bacillus subtilis* cloning

Antibiotic concentrations used in this study, unless otherwise indicated, were 100 µg/mL ampicillin, 1 µg/mL erythromycin, and 100 µg/mL spectinomycin. The full-length *B. subtilis tagA* gene was amplified from purified genomic DNA from *B. subtilis* 168 (*Bacillus* genetic stock center) and sub-cloned into the pBL113 shuttle vector (Table II) using *E. coli* XL10 (New England BioLab) to create pHisTagAFL. The hexahistidine-tag was incorporated during Gibson assembly (7). The truncated TagA construct (residues Met1-Val196) was constructed by amplifying the first 588 nucleotides of *tagA* and cloning into the pBL113 shuttle vector to create plasmid pHisTagAV196. *B. subtilis* was made competent as previously reported and transformed with 5-10 µL of pure plasmid to create strains RC168tagAFL and RC168tagAV196 (8). Homologous double-crossover at the *thrC* locus was verified using tryptophan/threonine auxotrophy and sequencing (Laragen Sequencing). The *tagA* gene was removed via allelic replacement using plasmid p*tagA::spec*, which was constructed by cloning 1 kb of DNA upstream and downstream of the *tagA* gene to flank the spectinomycin resistance cassette in the pIC56 plasmid; a portion of the spectinomycin cassette that was predicted to form a stem loop was removed to ensure that the native operon was not disrupted. Strain RC168tagAFL was made competent as previously described, transformed with an approximately 3 kb linear PCR product from p*tagA::spec*, and plated on LB agar plates with spectinomycin at 37°C to produce strain RC168tagAFLΔtagA. The *tagA* knockout was determined via colony PCR, and the DNA sequence was confirmed (Laragen Sequencing).

## SUPPLEMENTAL FIGURES

**Figure S1.**
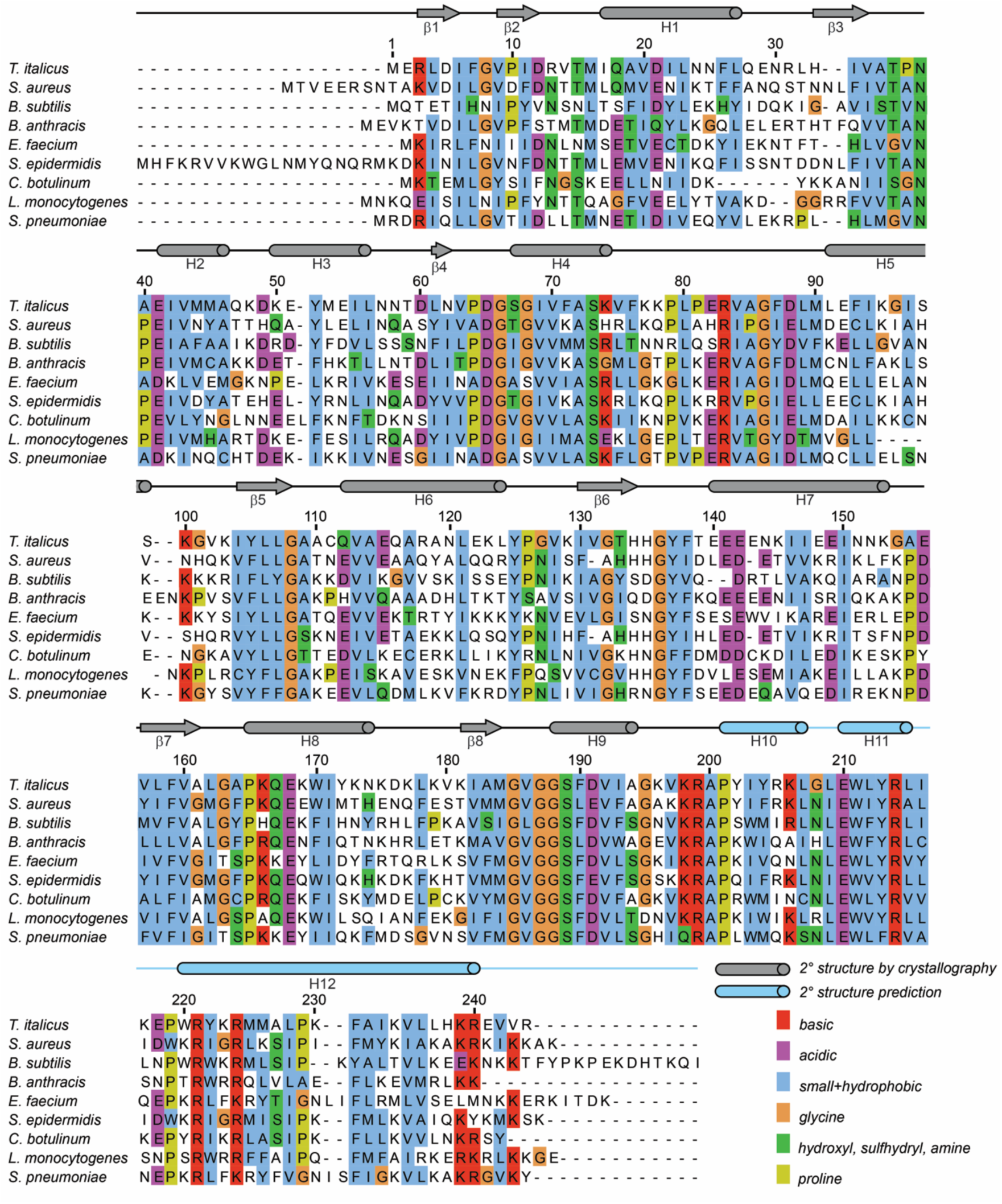
TagA primary sequence homology. The National Center for Biotechnology Institute’s Basic Local Alignment Tool (BLAST) was used to determine TagA sequence homologs with high sequence identity. The Clustal Omega multiple sequence alignment tool was used to generate a sequence alignment (9). Secondary structure is shown above the sequence and coloring is indicated in the key.

**Figure S2.**
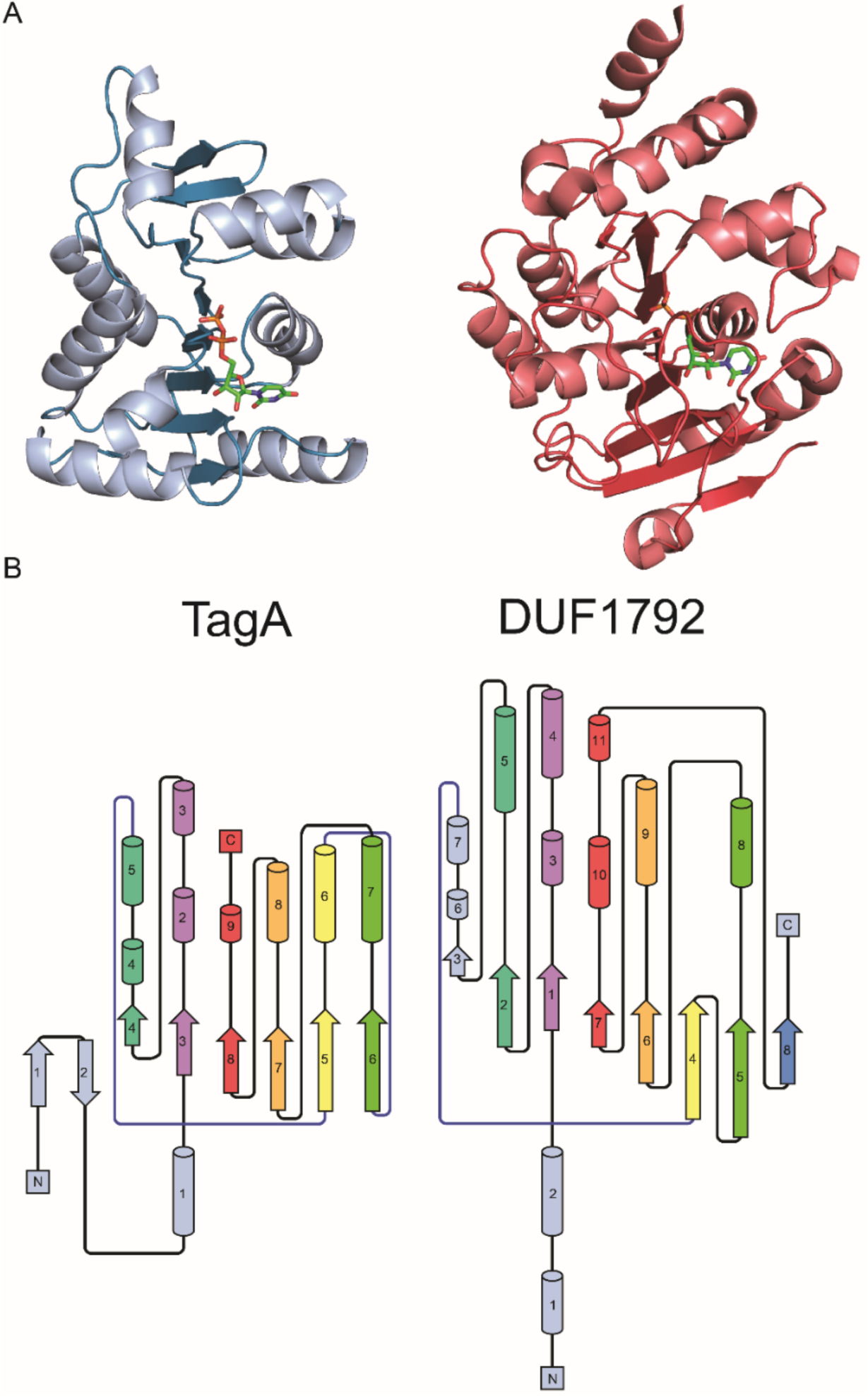
TagA structural comparison with DUF1792, a GT-D enzyme. (A) TagA aligns with DUF1792 with a DALI Z-score of 7.2 and an RMSD of 3.7 Å. Direct comparison reveals that a β-sheet composed of parallel β-strands is the main component of structural similarity. The tertiary organization of secondary structural elements between TarA and DUF1792 is significantly different. (B) Cartoon representation of secondary structure topology highlights that TagA has fewer β-strands in its sheet than DUF1792 and reinforces the dissimilarity in the order of secondary structural elements.

**Figure S3.**
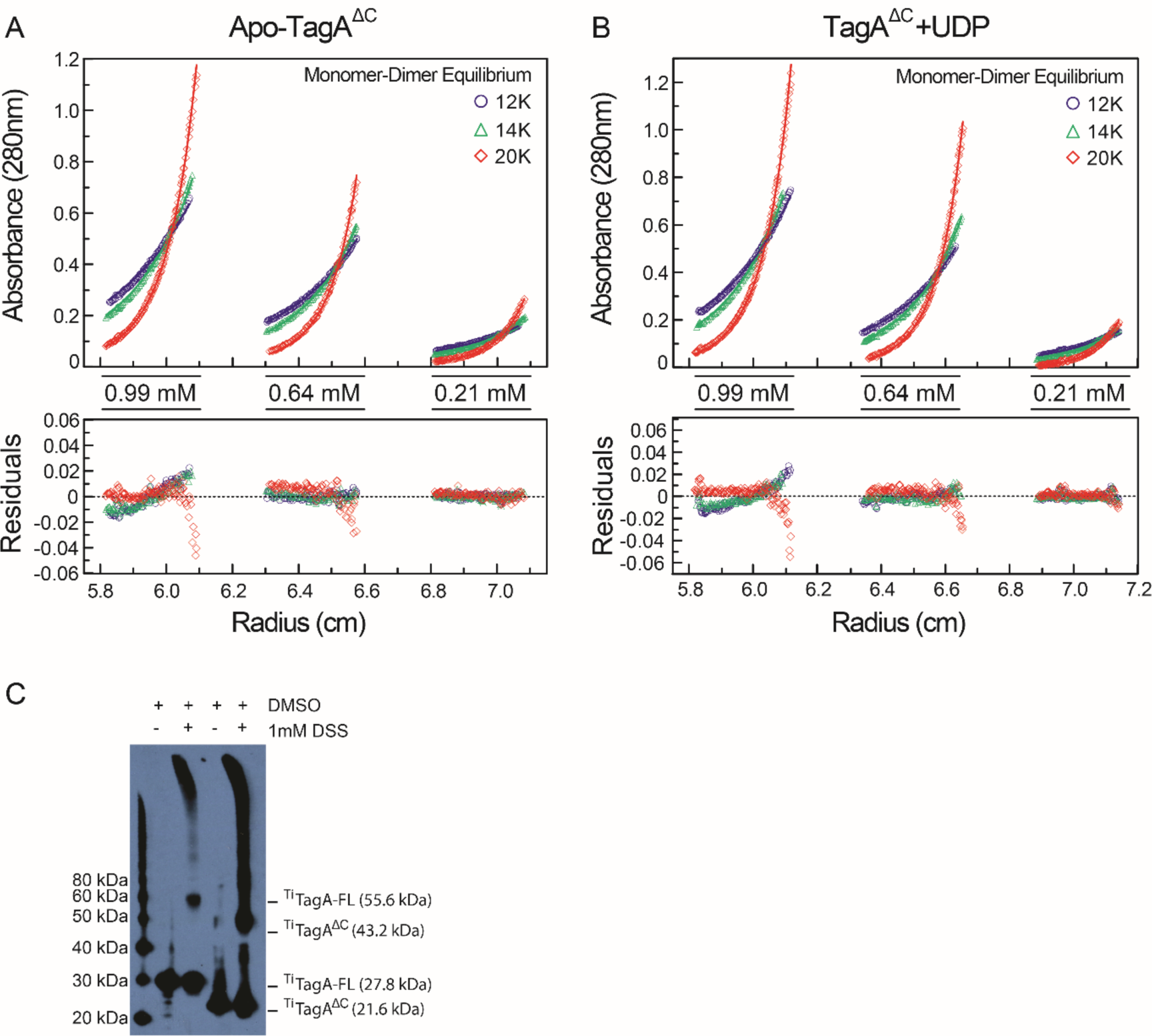
TagA oligomerization. The dissociation constant for TagA oligomerization was determined by equilibrium sedimentation analytical ultracentrifugation. The concentration distribution of TarA for three rotor speeds (12k, 14k, and 20k rpm) at three protein concentrations (0.99, 0.64, and 0.21 mM) for (A) apo-state TarA and (B) UDP-bound TagA. The lower panel shows the regression residuals for each protein concentration and centrifugal speed. The data were collected at 280 nm at 4°C and referenced against 50 mM Tris-HCl, pH 7.5, and 200 mM NaCl. (C) Crosslinking studies with disuccinimidyl suberate (DSS) in *E. coli* cells expressing *T. italicus* TagA constructs confirm that a dimer species is formed in the context of the cell. Both TagA and TagA^ΔC^ are monomeric under denaturing conditions (+ DMSO, - DSS); however, the addition of 1 mM DSS (+ DMSO, + DSS) produced a band corresponding to dimer species.

**Figure S4.**
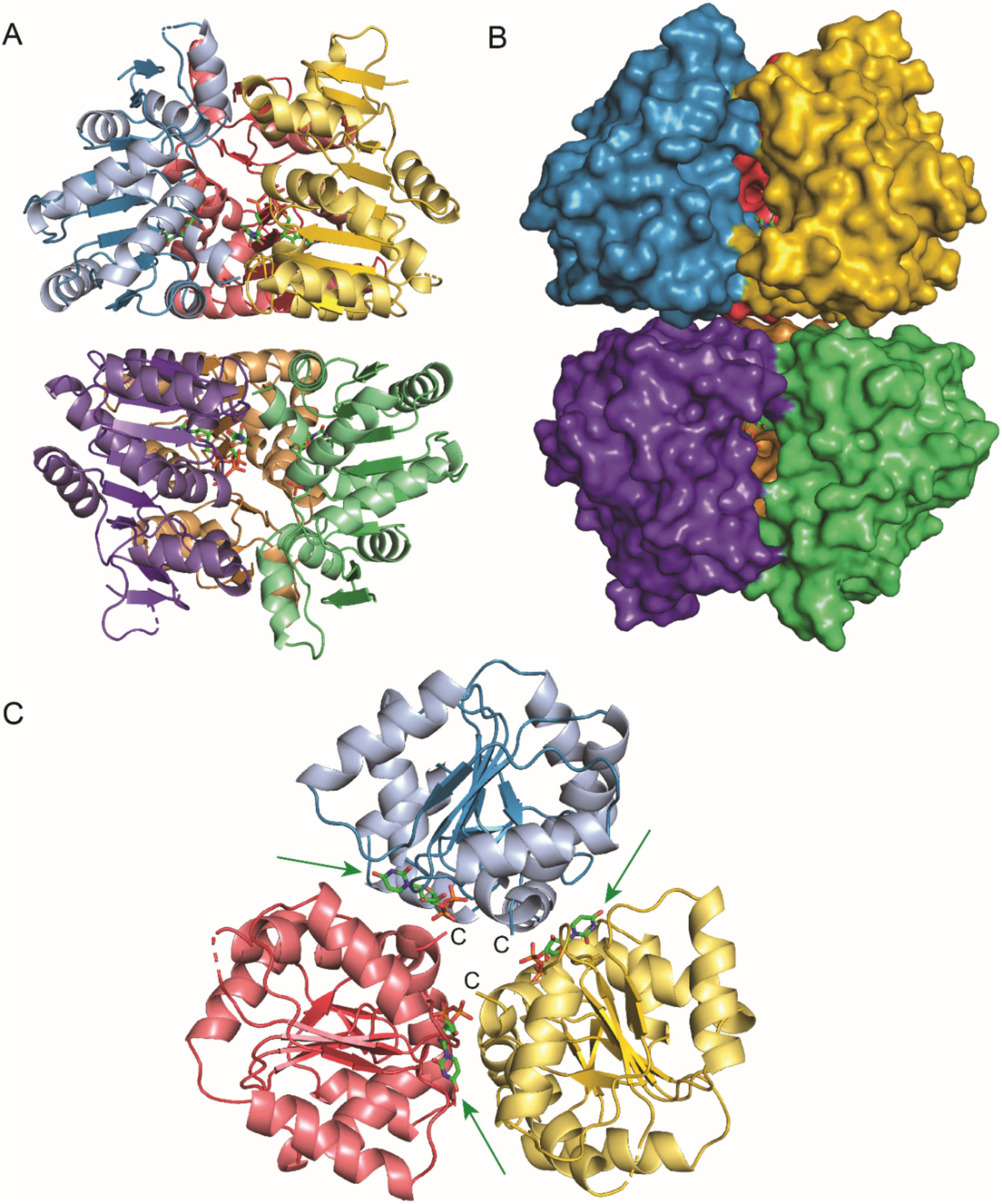
TagA^ΔC^:UDP complex crystallizes as a dimer of trimers. (A) Cartoon representation of the TagA^ΔC^:UDP complex. (B) Surface representation of the TagA^ΔC^:UDP complex. (C) View of one trimer unit within the crystallographic dimer. The C-termini are projected inward toward the center of symmetry. Green arrows indicate UDP, which can be seen at the interface between protomers of the trimer.

**Table S1.** – Strains and plasmids used in this study.

| Strain | Genotype/Description | Source |
| --- | --- | --- |
| RC168 | *B. subtilis* Wild type *trpC2* | BGSC |
| RC168tagAFL | RC168, *thrC*::pBL113-<sup>His</sup>*tagA* | Current study |
| RC168tagAFLΔtagA | RC168tagAFL, *thrC*::pBL113-<sup>His</sup>*tagA tagA::spec* |  |
| RC168tagAV196 | RC168 *thrC*::pBL113-<sup>His</sup>*TagA-V196* | Current study |
| RC21tagAFL | *E. coli* BL21(DE3) cells with pMAPLe4 containing *T. italicus* TagA | Current study |
| RC21tagAG195 | *E. coli* BL21(DE3) cells with pMAPLe4 containing *T. italicus* TagA (M1-G195) | Current study |
| Plasmid | Genotype/Description | Source |
| pMAPLe4 | *E. coli* plasmid that expresses proteins with N-terminal maltose binding protein fusion | (10) |
| pBL113 | *B. subtilis* – *E. coli* shuttle vector derived from pRDC19 (3) that integrates into the *thrC* locus in the *B. subtilis* genome, Amp<sup>R</sup>, Erm<sup>R</sup>, IPTG inducible | B. Lazazzera, UCLA (unpublished) |
| pHisTagAFL | *pBL113*-containing *B. subtilis tagA* gene with a 5' hexa-his tag. | Current study |
| pHisTagAV196 | *pBL113*-containing *B. subtilis tagA* gene with a 5' hexa-his tag truncated at G618 to produce a V196 translated product | Current study |
| pIC156 | *E. coli/B. subtilis* shuttle vector with spectinomycin<sup>R</sup> cassette. | (11) |
| *ptagA::spec* | pIC156 with genomic 1kb DNA flanking *tagA* from *B. subtilis* 168 inserted 5' and 3' to *spec*<sup>R</sup> cassette | Current study |

Author contributions: M.D.K., J.E.G., E.D.B., O.E.M. and R.T.C. designed research; M.D.K., J.E.G., O.E.M., D.C., M.R.S., M.P., G.K. and R.T.G. performed research; M.D.K., J.E.G., O.E.M., D.C., M.R.S., M.P., E.D.B. and R.T.C. analyzed data; M.D.K., J.E.G., E.D.B. and R.T.C. wrote the paper.

